# Metabolomic profiling reveals effects of marein on energy metabolism in HepG2 cells

**DOI:** 10.1101/176495

**Authors:** Baoping Jiang, Liang Le, Keping Hu, Lijia Xu, Peigen Xiao

## Abstract

Previous studies have suggested that *Coreopsis tinctoria* improves insulin resistance in rats fed with high-fat diet. But little is known about the antidiabetic effects of marein which is the main component of *C. tinctoria*. This study investigated the effects of ethyl acetate extract of *C. tinctoria* (AC) on insulin resistance (IR) in rats fed a high-fat diet. High glucose and fat conditions cause a significant increase in blood glucose, insulin, serum TC,TG and LDL-C, leading to an abnormal IR in rats. However, treatment with AC protects against HFD-induced IR by improving fasting serum glucose and lipid homeostasis. High glucose conditions cause a significant decrease in glycogen synthesis and increases PEPCK and G6Pase protein levels and Krebs-cycle-related enzymes levels, leading to an abnormal metabolic state in HepG2 Cells. However, treatment with Marein improves IR by increasing glucose uptake and glycogen synthesis and by downregulating PEPCK and G6Pase protein levels. The statistical analysis of HPLC/MS data demonstrates that Marein restores the normal metabolic state. The results show that AC ameliorates IR in rats and Marein has the potential effect in improving IR by ameliorating glucose metabolic disorders.

**Abbreviations:**

AC: ethyl acetate extract of *Coreopsis tinctoria*
TCA: Tricarboxylic acid
HepG2: hepatocellular carcinoma cell line
2-NBDG: 2-(N-(7-nitrobenz-2-oxa-1, 3-diazol-4-yl) amino)-2-deoxyglucose
G6Pase: glucose-6-phosphatase
PEPCK: phosphoenolpyruvate carboxykinase
IR: insulin resistance
HFD: high-fat diet
SDHA: succinate dehydrogenase flavoprotein subunit
ACO2: aconitase 2
IDH2: isocitrate dehydrogenase 2
CS: citrate synthase
FH: fumarate hydratase
MDH2: malate dehydrogenase
DLST: dihydrolipoamide S-succinyltransferase

## Introduction

Type 2 diabetes (T2DM) is a progressive disease characterized by deterioration of glycaemia and escalating therapeutic complexity ^[1]^. In diabetes mellitus, chronic hyperglycemia develops as a consequence of decreased insulin action from impaired insulin secretion and insulin resistance (IR) ^[2, 3]^. Once chronic hyperglycemia is established, it in turn aggravates IR, forming a vicious cycle that is collectively called glucose toxicity^[4, 5]^. The liver is the primary organ responsible for regulating glucose homeostasis ^[6]^. Hepatic IR leads to altered glucose metabolism and hyperglycemia, which is characterized by the inability of insulin to inhibit hepatic gluconeogenesis by suppressing unidirectional enzymes, namely, phosphoenolpyruvate carboxykinase (PEPCK) and glucose 6-phosphatase (G-6Pase)^[7, 8]^. G-6Pase plays a role in glucose homeostasis ^[9]^. PEPCK is a key rate-limiting enzyme of gluconeogenesis. The activities of G-6Pase and PEPCK are increased significantly in the liver of diabetic rats ^[8]^. Inhibition of G-6Pase and PEPCK enzymes may interfere with gluconeogenesis and can be useful in treating diabetic hyperglycemia ^[10-13]^. In the liver mitochondrial PEPCK (PEPCK-M)adjoins its profusely studied cytosolic isoform (PEPCK-C) potentiating gluconeogenesis and TCA flux, so hepatic PEPCK is required to sustain the Krebs cycle ^[14]^. In addition, the Krebs cycle completes the oxidation of glucose and plays an important role in the glucose metabolism. Therefore, methods that enable simultaneous measurement of numerous cellular metabolic intermediates in gluconeogenesis and the Krebs cycle are required to elucidate the mechanisms of glucose metabolism disorders.

Metabolomics is a top-down systems biology approach whereby metabolic responses to biological interventions or environmental factors are analyzed and modeled ^[15]^. Metabolomics measures perturbations in metabolites reflecting changes of metabolism caused by environmental factors, and provides insights into the global metabolic status of the entire organism by monitoring the entire pattern of low molecular weight compounds rather than focusing on an individual metabolic intermediate ^[16]^. Deregulations of metabolic processes are expected to be directly or related with relevant disease end-points, which are represented by the levels of metabolites ^[17]^. Recent longitudinal metabolomic studies have found correlations between circulating metabolites and prediabetes, future development of IR, or type 2 diabetes in humans. For example, increasing in circulating aromatic amino acids (AAAs) and branched-chainamino acids (BCAAs) are biomarkers of risk ^[9, 18, 19]^.

*Coreopsis tinctoria* Nutt. (Asteraceae), a traditionally used preparation for diabetes treatment in Portugal ^[20, 21]^, is a plant native to North America that has spread worldwide. Our previous study showed that *C. tinctoria* increases insulin sensitivity and regulates hepatic metabolism in rats fed a high-fat diet ^[22]^.Since these activities are closely related to metabolic regulation and IR, we tried to identify the protective effect of ethyl acetate extract of *C. tinctoria* Nutt (AC) on IR, and its possible mechanism of action. Chalcones (okanin and butein derivatives) are the main constituents of ethyl acetate extract of *C. tinctoria* and among them, identified Marein (okanin-4’-O-β-D-glucopyranoside) as the main metabolite ^[23]^. Marein has many beneficial biological activities, including antihyperlipidemic ^[24]^, antioxidative ^[25]^, antidiabetic ^[26]^, and antihypertensive effects ^[27]^. Previous research has also found that Marein prevented tert-Butyl-Hydroperoxide and cytokine induction in a mouse insulinoma cell line (MIN6) through the inhibition of the apoptotic signaling cascade^[20]^. In addition, Marein promotes pancreatic function recovery in streptozotocin-induced glucose-intolerant rats ^[26, 28]^. Our previous studies found that Marein could be the main active compounds of *C. tinctoria* in improving IR^[29]^. The protective effects of marein on high glucose-induced metabolic disorder via IRS1/AKT/AS160 signal pathway in HepG2 cells^[30]^. These findings focused on *in vivo* anti-diabetic effect led us to further investigate the underlying mechanism, as to our knowledge no studies on the mechanisms of metabolic disorder of Marein have been reported, especially on energy metabolic level.

In this study, we employed LC/MS-based metabolomics techniques to explore the potential targets and mechanisms of Marein for the attenuating IR and type 2 diabetes. Our study revealed involvement of glucose metabolism and the Krebs cycle in the protective effects of Marein in a high-glucose and fat-induced IR model.

## Materials and methods

### Materials

The human liver hepatocellular carcinoma cell line HepG2 was purchased from the Cell Bank of the Chinese Academy of Sciences (Beijing). Dulbecco’s modified Eagle’s medium (DMEM), fetalbovine serum (FBS), and other tissue culture reagents were purchased from Gibco (Life Technologies, USA). Other reagents were purchased from Sigma-Aldrich Co. (St. Louis, MO, USA) unless otherwise indicated. AC was analyzed by HPLC (Waters 2690) with a diode-array detector (Waters 2487) scanning from 200–600 nm. Ethyl acetate extract was seperated by Shim-pack VP-ODS column (150 × 4.6 mm, 5 μm) with the optimum condition as described previously^[28, 31]^. Marein was extracted from *C. tinctoria Nutt.* by our lab. And we also purchased the compound Marein from Chromadex (00013126-604, California, USA), and the purities exceeded 99%. Deionized water was used in all experiments. All of the other chemicals and reagents were of analytical grade. The following antibodies were used in immunoblotting experiments and at the indicated dilutions: rabbit polyclonal anti-PCK1 (PEPCK) (ab28455, 1:1000), rabbit polyclonal anti-G6Pase (ab83690, 1:1000), rabbit monoclonal anti-aconitase 2 (ACO2) (ab129069, 1:10000), rabbit monoclonal anti-malate dehydrogenase (MDH2) (ab181857, 1:10000), mouse monoclonal anti-isocitrate dehydrogenase 2 (IDH2) (ab184196, 1:1000), mouse monoclonal anti-fumarate hydratase (FH) (ab113963, 1:50000), rabbit polyclonal anti-citrate synthase (CS) (ab96600, 1:1000), mouse monoclonal anti-succinate dehydrogenase flavoprotein subunit (SDHA) (ab14715, 1:10000), rabbit monoclonal anti-SDHB (ab178423, 1:5000, abcam, UK), and anti-β-actin antibody (1:1000, CW0096A, CWBio). Dihydrolipoamide S-succinyltransferase (DLST)

### Animals and drug treatments

Sprague Dawley rats were randomly divided into six groups of 10 animals each. The control group that was given deionized distilled water to drink and fed standard rat chow 32 composed of 60% vegetable starch, 12% fat, and 28% protein. The high-fat diet (HFD) model group^[32]^ that was given deionized distilled water and fed a high fat diet of 60% fat, 14% protein and 26% carbohydrate. The metformin group was administrated with 200 mg/kg of metformin as a positive control by oral gavage and fed a HFD. The rats in the AC three groups were as follows: low dose of AC (150 mg/kg of body weight + HFD), middle dose of AC (300 mg/kg of body weight + HFD), high dose of AC (600 mg/kg of body weight + HFD). The study was approved by the Ethics Committee of the Institute of Medicinal Plant Development, CAMS&PUMC (Beijing, China). All experimental procedures were performed in accordance with relevant guidelines approved by the Ethics Committee of the Institute of Medicinal Plant Development, CAMS&PUMC.

### Cell culture and drug treatment

HepG2 cells were cultured in low glucose DMEM (5.5 mmol/L glucose) that was supplemented with 10% FBS and 1% antibiotics at 37°C in humidified air containing 5% CO_2_. Cells in the exponential phase of growth were used in all of the experiments. HepG2 cells were grown for 3 days and then divided into different groups for the treatments. The treatment groups (Marein, dissolved in DMSO, not more than 0.5 percent) were as follows: (1) Control: incubated in low glucose DMEM for 72 h; (2) High glucose treatment: incubated in DMEM containing 55 mmol/L glucose for 72 h; (3-6) Marein and glucose treatment: incubated in DMEM containing Marein (40, 20, 10, or 5 μ mol/L) or 0.5 mmol/L metformin (Positive control) for 24 h and then incubated in DMEM containing 55 mmol/L glucose for 72 h. 100 nmol/L insulin was added into the all treated group for 30 min before all experiments.

### Detection of glucose uptake

Glucose uptake rate was measured by adding 2-(N-(7-nitrobenz-2-oxa-1, 3-diazol-4-yl) amino)-2-deoxyglucose, a fluorescently labeled deoxyglucose analog (2-NBDG, Cayman Chemical), as a tracer to the culture medium as previously reported ^[33]^. 2-NBDG uptake was then measured after stimulating the cells for 15 min with 1×10^−7^ mol/L insulin as our previous described^31^. The cells treated by drugs were then washed twice and incubated with 100 μmol/L of 2-NBDG in glucose-free culture medium for 20 min. The cells cultured in medium without 2-NBDG were used as a negative control. The cells were washed twice prior to fluorescence detection using a microplate reader (Infinite 1000 M, Tecan, AUSTRIA) with fluorescence excitation at 488 nm and emission detected at 520 nm.

### Determination of ATP, pyruvate and lactate concentrations

The cellular ATP content was detected by using an ATP content kit (S0026, Beyotime Institute of Biotechnology, China). Briefly, the cell pellets were collected by centrifugation at 1000 ×*g* for 5 minutes after digestion with 0.25% trypsin. The homogenate was incubated in boiling water for 30 minutes, and the supernatant was aspirated after centrifugation at 5000 ×*g* for 10 minutes. An ATP assay kit was used for all of the samples. The results were analyzed with a luminometer (Fluoroskan Ascent FL, USA).

The cellular lactate and pyruvate concentration was detected by using a lactate assay (A019-2, Nan Jing Jian Cheng Bioengineering Institute, Nan Jing City, China) and pyruvate assay kit (A081, Nan Jing Jian Cheng Bioengineering Institute, Nan Jing City, China) respectively. Briefly, cell pellets were collected by centrifugation at 500 ×*g* for 10 minutes after scraping the cells with a cell scraper. Cells were washed twice with PBS, then resuspended in PBS and lysed with a sonicator. The homogenate was detected using a UV/Visible Spectrophotometer (Lambda 35, Perkin Elmer, USA).

### RNA extraction and quantitative real-time PCR (qPCR)

Cells treated with high glucose or Marein were then collected to isolate total RNA. Total RNA was reverse transcribed, and *PEPCK, G6Pase, FH, MDH2, SDHA, SDHB, IDH2, ACO2, CS* and *GAPDH* gene expression was measured by QPCR using iCycler thermocycler (BioRad, Hercules, CA) as previously described ^[34]^. The primers of these genes are listed in the Table 1. All samples were analyzed in triplicate, and the gene expression levels were normalized to control human *GAPDH* values. The relative expression of treated samples was normalized to that of untreated cells according to ΔCt analysis.

### Western blot analysis

All immunoblots were of the cell lysate with 1×10^7^ cells per sample with RIPA cell lysis buffer (50 mM Tris-HCl, pH 7.4, 1% NP-40, 0.5% Na-deoxycholate, 0.1% SDS, 150 mM NaCl, 2 mM EDTA and 50 mM NaF). PEPCK and G6Pase were detected with anti-PEPCK and anti-G6Pase antibodies respectively, both used at a dilution of 1:1000. FH, MDH2, SDHA, SDHB, ACO2, CS, IDH2 and GAPDH were detected by anti-FH, anti-MDH2, anti-SDHA, anti-SDHB, anti-ACO2, anti-CS, anti-IDH2 and anti-GAPDH for 4h following the instructions respectively. The secondary antibodies goat anti-mouse IgG and goat anti-rabbit IgG were peroxidase-conjugated at a dilution of 1:10000 and 1:20000 respectively. Immune complexes were visualized with ECL chemiluminescence (GE Healthcare, Little Chalfont, UK). Densitometric analyses were performed with ImageJ2 software for Microsoft Windows.

### Metabolite profiling

Metabolite profiling was performed as previously reported ^[35]^. Targeted metabolomic experiment was analyzed by TSQ Quantiva (Thermo, CA). C18 based reverse phase chromatography was utilized with 10mM tributylamine, 15mM acetic acid in water (pH ∼6)and 100% methanol as mobile phase A and B respectively. This analysis focused on tricarboxylic acid cycle, glycolysis pathway, pentose phosphate pathway, amino acids and purine metabolism. In this experiment, we used a 25-minute gradient from 5% to 90% mobile B. Positive-negative ion switching mode was performed for data acquisition. Cycle time was set as 1 second and a total of 138 ion pairs were included. The information of partial ion pairs are listed in Table 2. The resolution for quartile 1 and quartile 3 are both 0.7FWHM. The source voltage was 3500v for positive and 2500v for negative ion mode. Sweep gas was turned on at 1(arb) flow rate. In the experiments, metabolites were extracted from 1× 10^7^ HepG2 cells and finally redissolved in 80 μl of 80% methanol. One microliter of sample was loaded for targeted metabolomics analysis. We used specific sample as quality control (QC) instead of internal standard. Total ion current and chromatographic patterns were evaluated. And Table 3 shows three representative compounds in QC runs.

### Statistical analysis

The MS raw data of cellular extracts was processed using Thermo Xcalibur (Thermo, USA). Analyzed spectral data was transformed to contain aligned peak area with the same mass/retention time pair as well as normalized peak intensities and sample name. The main parameters for data processing were set as follows: mass range (80-1200 Da), mass tolerance (5 ppm), intensity threshold (100 counts), retention time (0-20 min) and retention time tolerance (0.2 min). The resulting data were mean-centered and pareto-scaled prior to statistical analysis by Principal Component Analysis (PCA) to differentiate each group. PCA was used to visualize the maximal difference of global metabolic profiles^[36]^. All data were expressed as means±standard error of the mean (SEM). Statistical significance was calculated with one-way ANOVAs and post hoc Turkey’s tests in GraphPad Prism 5.0 (GraphPad Software, San Diego, CA, USA).. Differences were considered to be statistically significant when P<0.05. All experiments were performed at least four times.

## Results

### AC partially improves serum glucose and lipid homeostasis and protects against HFD-induced IR in HFD-fed rats

To explore the effect of AC in rats, we constructed the animal IR model induced by high glucose and fat diet. *The results showed that* the serum total cholesterol (TC) and triglyceride (TG) concentrations were lower than those in the chow diet-fed animals at the end of the 8-week HFD (Fig. 1A and 1B). The analysis of serum lipoproteins revealed that the hypercholesterolemia in rats fed a HFD was associated with increased low-density lipoprotein-cholesterol (LDL-C) and decreased high-density lipoprotein-cholesterol (HDL-C), typical phenotypes of diabetic dyslipidemia (Fig. 1C and 1D). Treatment with AC for 8 weeks significantly improved the lipid profiles and reduced the serum levels of glucose and insulin. AC or metformin (MET; 200 mg/kg) significantly reduced the fasting glucose and insulin levels (Fig. 1E and 1F). Moreover, treatment with Marein increased insulin sensitivity index and improved the insulin tolerance (Fig. 1G and 1H). Although, HFD rats showed a clear increase in body weight, there are no significantly changes in drug treated group except the positive control compared with HFD group. Histological examination of the liver in HFD-treated rats showed lipid accumulation and fatty degeneration in the hepatocytes; however, treatment with 100 mg/kg marein significantly attenuated the formation of fat vacuoles in the liver sections (Fig. 2).

**Figure 1.**
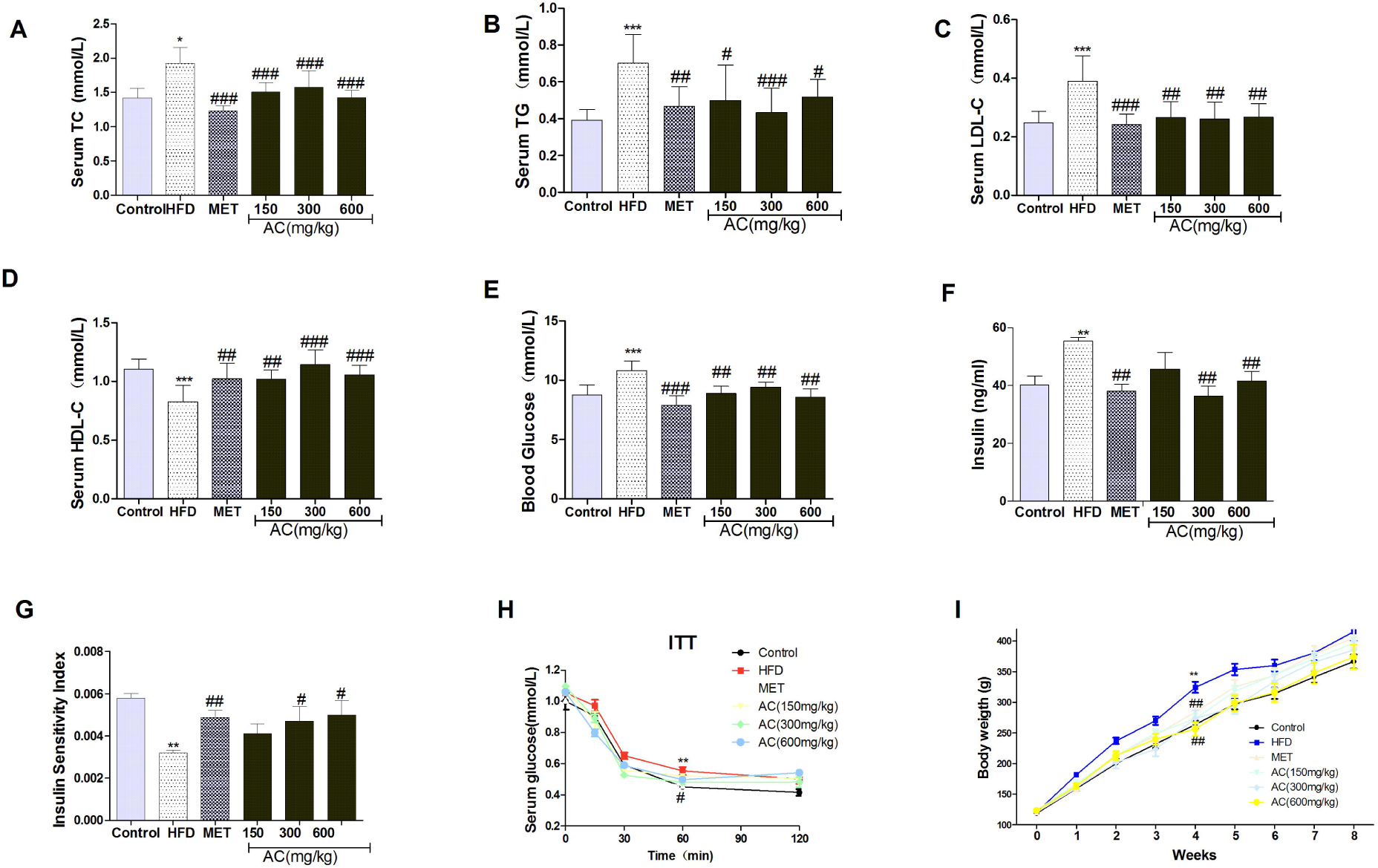
AC improves serum glucose and lipid homeostasis and protects against HFD-induced IR. The serum levels of TC (A), TG (B), LDL-C (C), HDL-C (D), fasting blood glucose (E), fasting insulin (F) were measured. G, Insulin sensitivity index [ISI =1/(fasting insulin × fasting plasma glucose)]. H, insulin tolerance tests (ITT). I, body weight Values are the means ± SEM (n = 10). .* *P<0.05* vs the control group; ** *P<0.01* vs the control group; # *P<0.05* vs HFD-treated group; ## *P<0.01* vs HFD-treated group.

**Figure 2.**
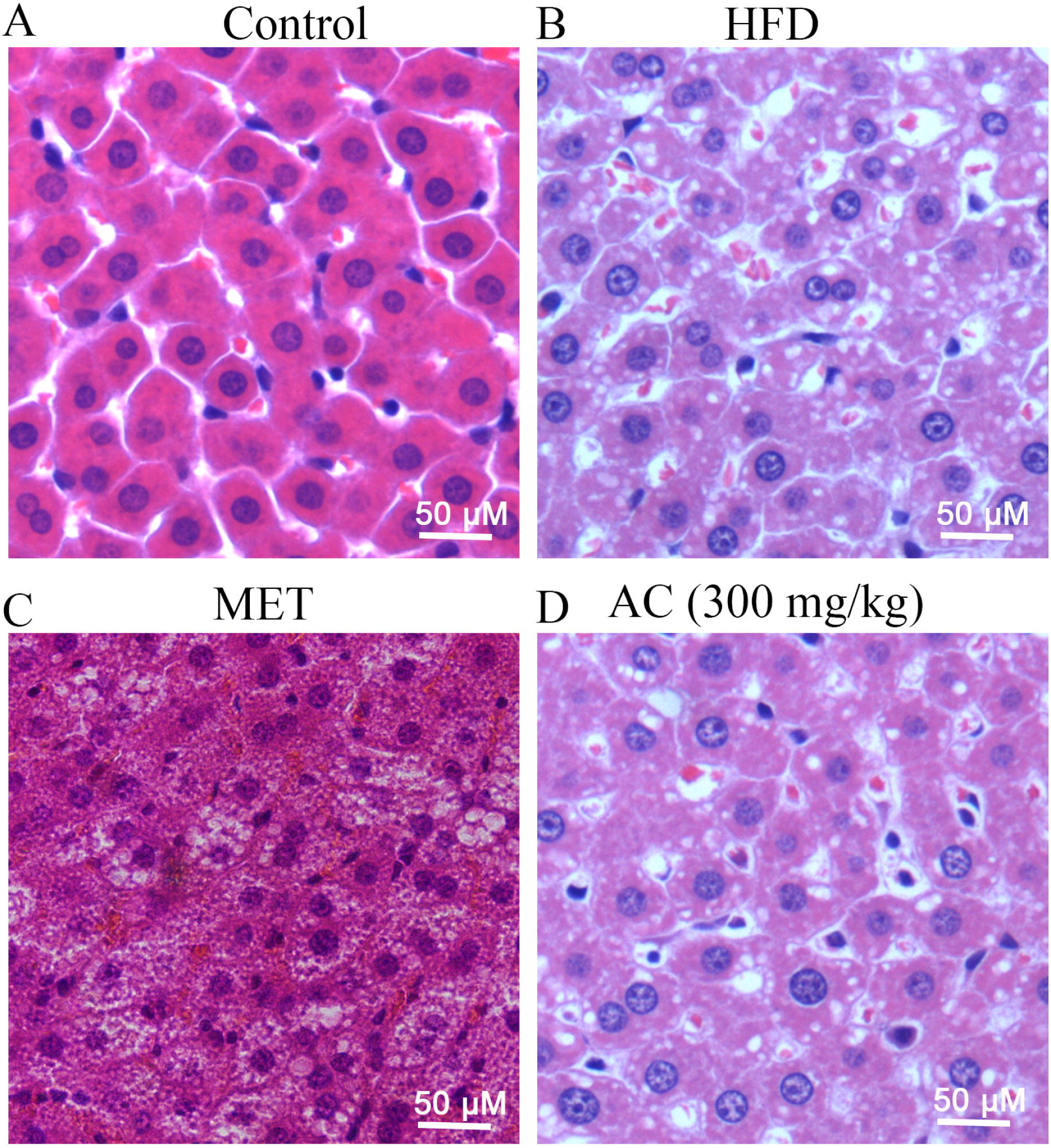
Effects of AC on lipid accumulation and steatosis. Representative hematoxylin and eosin staining of the liver (magnifications, × 200).

### Marein increases glucose uptake, hexokinase activity, and glycogen synthesis and decreases RNA and protein expression of PEPCK and G6Pase in high-glucose-induced IR model cells

We evaluate the effect of marein (Fig. s1) extracted from *C. tinctoria* on glucose consumption. 55 mmol/L glucose-induced cell model of IR was successfully constructed with HepG2 cells. Glucose decreased 2-NBDG uptake in a both time-dependent and dose-dependent manner (Fig. 3A). The results showed that Marein had a potential effect on promoting the uptake of glucose into HepG2 cells. The dose-dependent effects of Marein treatment for 24h were investigated from 1.25 μmol/L to 40 μmol/L. After treatment with 5 μmol/L of Marein, the 2-NBDG uptake into HepG2 cells reached the highest level, which was the plateau level (Fig. 3B). Then, the effect of Marein on IR induced by high glucose was investigated with 2-NBDG. Pre-treatment of HepG2 cells for 24h with all concentrations of Marein obviously increased glucose uptake that approach to 90.4% of control with 10 μmol/Lol/L Marein (Fig. 3C), although no significant difference of cell survivals were observed in HepG2 cells treatment with Marein (Fig. 3D),. We thus selected 10 and 5 μmol/L Marein to use in the subsequent experiments. The IR model with HepG2 cells (55 mmol/L glucose) causes a decrease in the glycogen content and hexokinase (a key enzyme in glycogen synthesis) activity. Both concentrations (5 and 10μmol/L Marein) inhibited the effect of 55 mmol/L glucose (Fig. 3E and 3F). Marein can thus increase hexokinase activity and glycogen synthesis in a dose-dependent manner in an IR model where these were previously suppressed.

**Figure 3.**
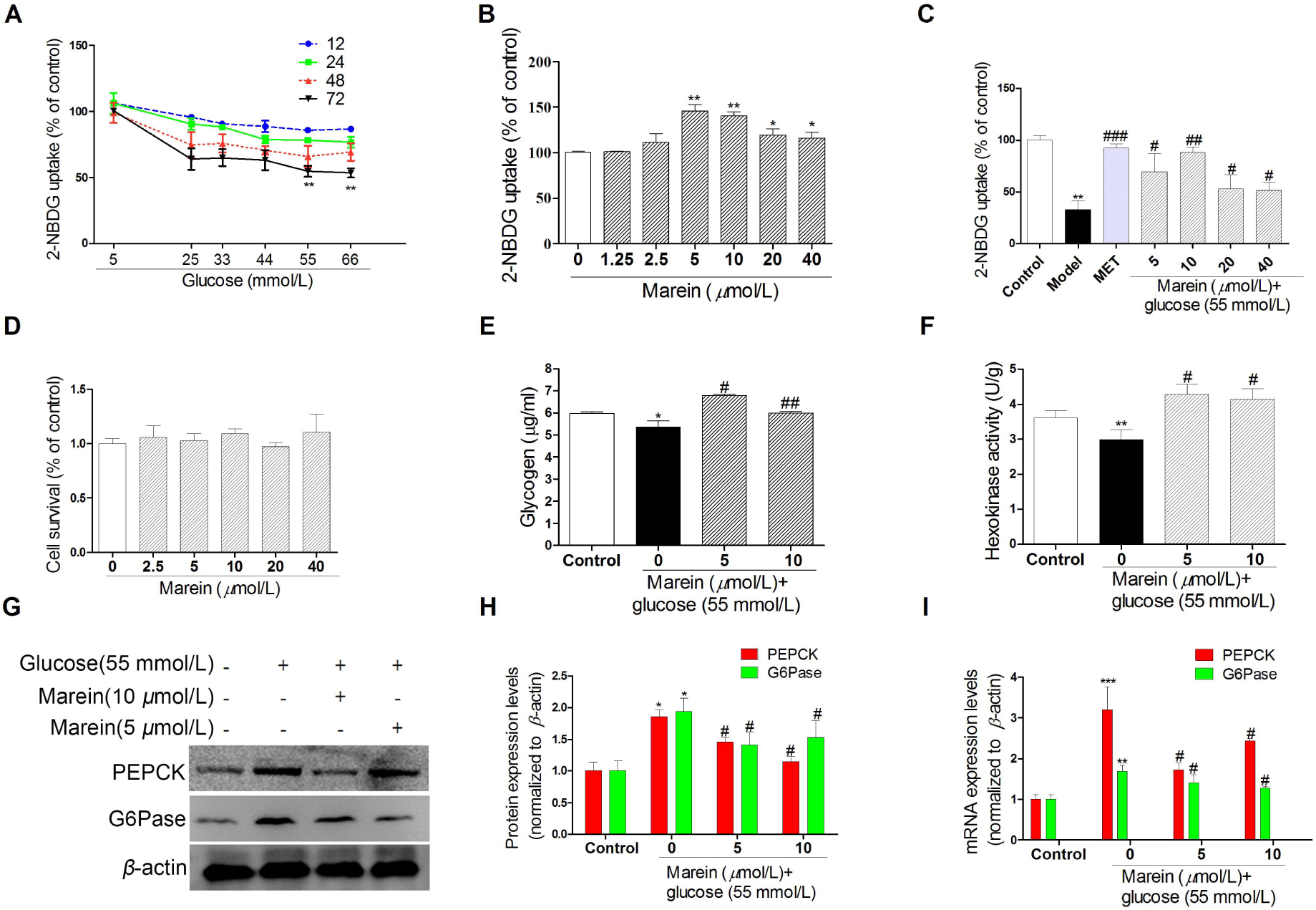
Marein inhibits the decrease of glucose uptake and the imbalance of glucose metabolism induced by high glucose in HepG2 cells. (A) Dose-dependent and time-dependent effect of glucose on 2-NBDG uptake. (B) Dose-dependent effect of Marein on 2-NBDG uptake. HepG2 cells were incubated for 24h. (C) Glucose uptake expressed as a percent of control group are means ± SD of at least 4 different samples per condition. (D) Cell viability shows the toxicological effect of Marein in HepG2 cells. (E) Effect of Marein on glycogen content. (F) Effect of Marein on Hexokinase activity. (G) Bands of representative experiments for PEPCK and G6Pase. (H) Densitometric quantification of PEPCK and G6Pase. (I) Effect of Marein on mRNA expression of PEPCK and G6Pase. All experiments were performed at least 4 times. * *P<0.05* vs the control group; ** *P<0.01* vs the control group;*** *P<0.001* vs the control group; # *P<0.05*vs 55 mmol/L glucose-treated group; ## *P<0.01* vs 55 mmol/L glucose-treated group.

In the hepatocyte, PEPCK and G6Pase levels and gluconeogenesis are stimulated by inhibition of AKT and are restrained by the synthesis of glycogen in a situation of IR ^[37]^. In this study, 55 mmol/L glucose caused an increase in protein and mRNA levels of PEPCK and G6Pase, but Marein inhibited these alterations (Fig. 3G-I).

### Multivariate statistical analysis of HPLC/MS data summarizes metabolic differences in treatment groups

Measured levels of small molecule metabolites of enzymes involved in glucose metabolism and the Krebs cycle are used to verify the changes of these enzymes. Representative HPLC/MS data of cell samples from the high glucose treated cells, control cells and Marein-treated group were used in statistical analysis, as shown in Fig. 4. The three-dimensional PCA score plot for the first three principal components (PC1, PC2 and PC3) with clustering of each group. The clear separation of the high glucose group and control group suggests that severe metabolic disturbance occurs in the IR model cells (Fig. 4A, 4B). Marein (5 μmol/L and 10 μmol/L) group was clearly separated from the model group (high glucose) and partially overlapped with control group, suggesting Marein had apparent effect on the IR. The PCA score plots of PC1, PC2 and PC3 revealed that data points, each representing one sample, were clustered in a way that allowed Model group (high glucose) to be clearly separated from control, Marein (10 μmol/L), Marein (5 μmol/L) and positive control groups along PC1 (Fig. 4C, 4D, 4E). Changes in important positions in a network more strongly impact the pathway than changes occurring in marginal or relatively isolated positions. MetaboAnalyst 3.0 revealed that differential metabolite content is important for the normal response to high glucose group, and multiple pathways are altered in high glucose and Marein group (Fig. 4F). The impact-value threshold calculated via pathway topology analysis was set to 0.10 ^[38]^, and 27 of the regulated pathways were identified as potential target pathways (Fig. 4F and Table 4) of marein in high glucose-induced model.

**Figure 4.**
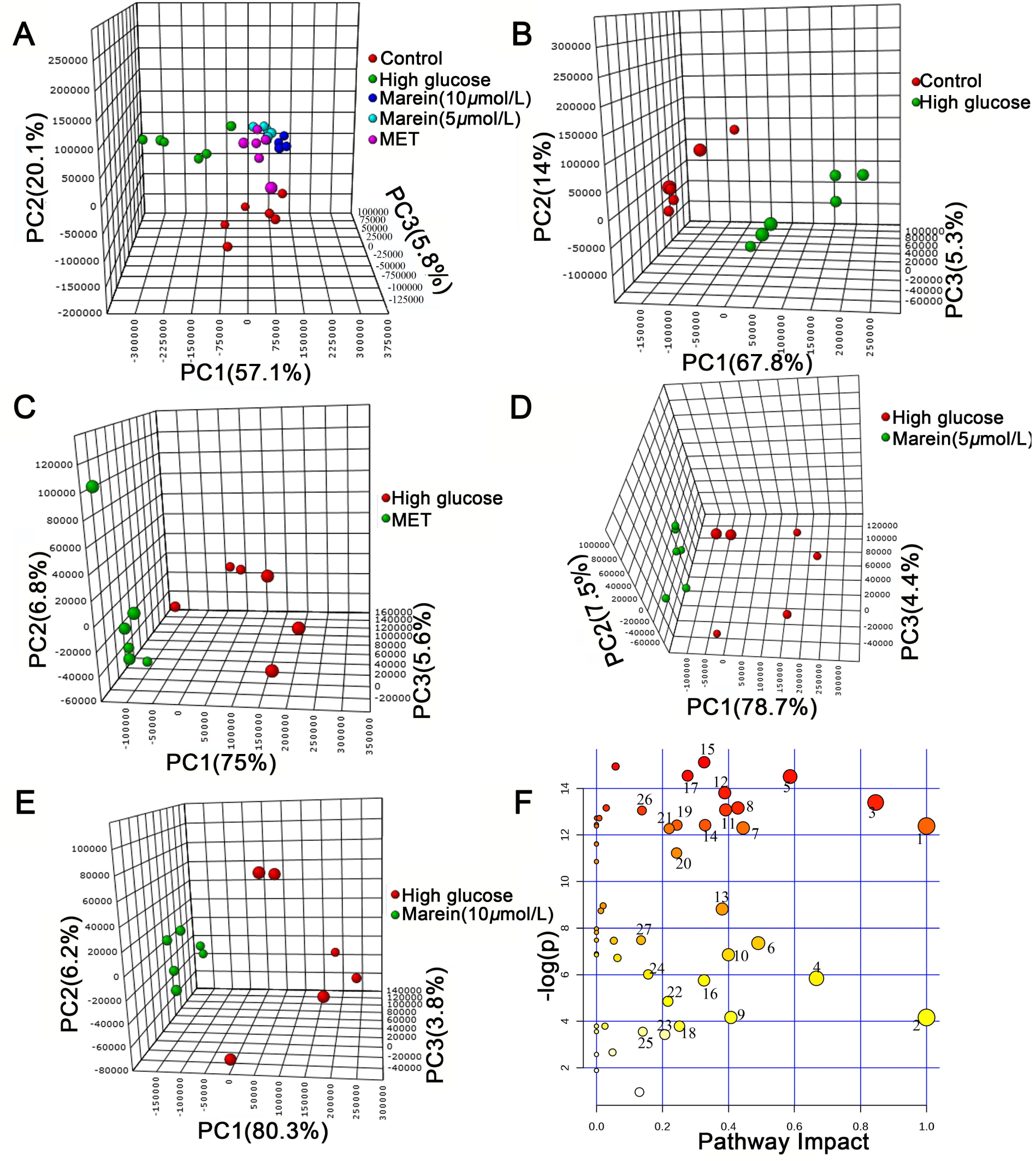
PCA scores plots for HPLC/MS data of all groups. (A) The three-dimensional PCA score plot of metabolic states of normal control group (red), high glucose group (green •), 10 μmol/L Marein group (blue •), 5 μmol/L Marein group (bright blue •) and Positive control group (pink •). (B) The three-dimensional PCA score plot of metabolic states of normal control group (red •) and high glucose group (green •). (C) The three-dimensional PCA score plot of metabolic states of high glucose group (red •) and Positive control group (green •). (D) The three-dimensional PCA score plot of metabolic state of high glucose group (red •) and 5 μmol/L Marein group (green •). (E) The three-dimensional PCA score plot of metabolic state of high glucose group (red •) and 10 μmol/L Marein group (green •). (F) Summary of pathway analysis with MetPA. Positive control is 0.5 mmol/L metformin.

Furthermore, unsupervised hierarchal cluster analysis revealed the fluctuation of levels across different groups, as visualized by a heat map (Fig. 5A). Metabolic substrates of different treatment groups were changed in different degrees. Z-score plots were constructed to identify metabolic changes distinct between high glucose group and control group. 137 metabolites exclusive to cells after treatment with high glucose (red plot) were identified. However, treatment with 10 μmol/L Marein significantly reduced the levels of these metabolites (green plot, Fig. 5B).

**Figure 5.**
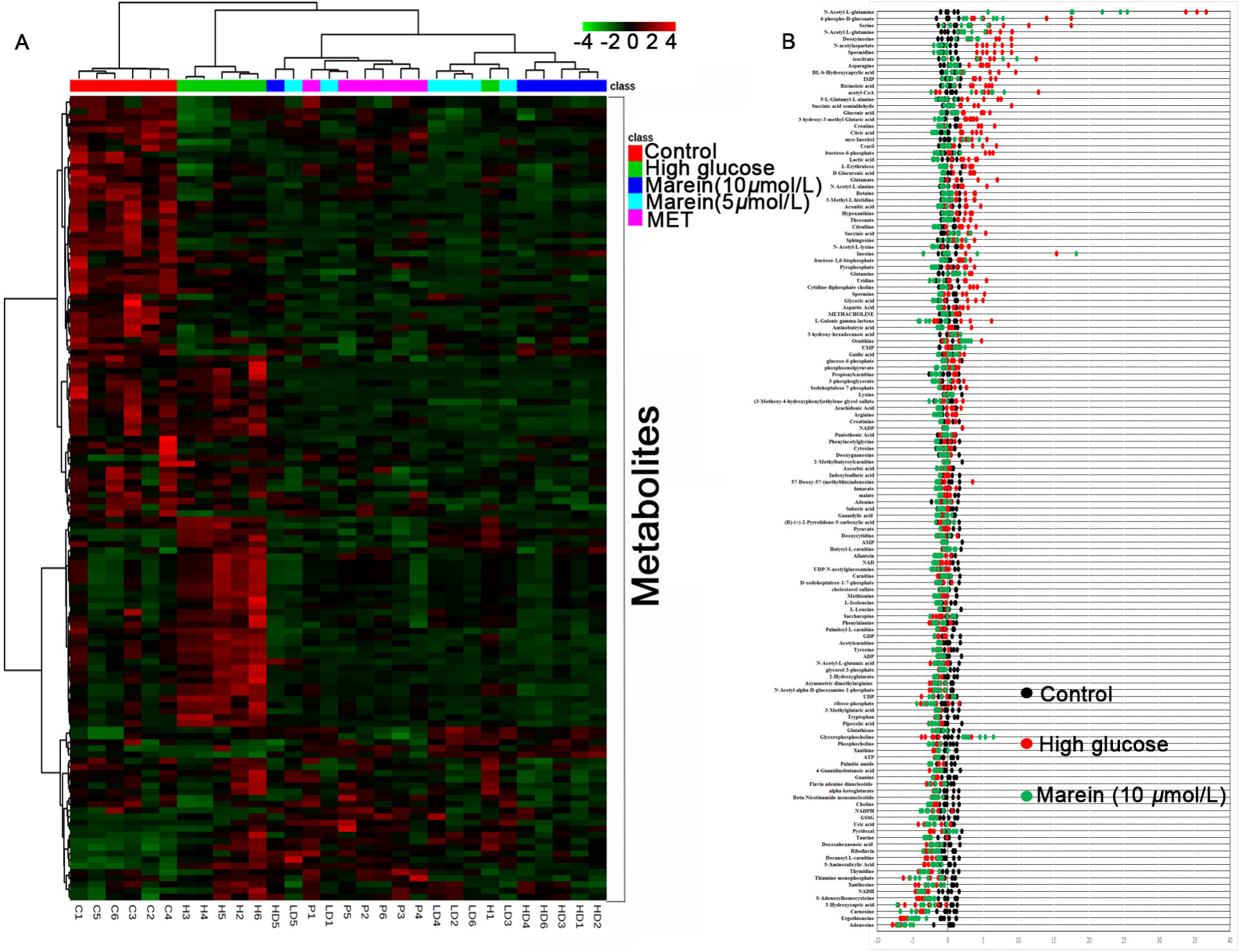
The variance analysis of metabolites based on the data of HPLC/MS of cells. (A) Heat map visualization of the correlation analysis of all metabolites. Row: groups (red: normal control; green: high glucose; blue: 10 μmol/L Marein; bright blue: 5 μmol/L Marein; pink: Positive control); columns: metabolites; Cell color key indicates the cluster score, green: Lowest, red: highest. (B) High glucose-based z-score plot of named metabolites for comparison between normal control (black), high glucose (red) and 10 μmol/L Marein (green). Data are shown as standard deviation from the mean of respective sham. Each dot represents a single metabolite in 1 sample; n=4 per group.

### Effects of Marein on energy metabolism intermediates in high glucose-induced IR model cells

High glucose-treated cells exhibited changes in the levels of metabolic intermediates that participate in energy metabolism, including the glycolytic pathway and the Krebs cycle. Concentrations of major intermediates involved in glycolysis, such as dihydroxyacetone phosphate (DHAP; 0.48-fold, p= 8×10^−3^), and 3-phosphoglycerate (0.61-fold, p= 0.04), were significantly decreased in the high glucose group, compared with control group. However, 10 μmol/L Marein reduced the effect of high glucose on these two metabolites (Fig. 6).

**Figure 6.**
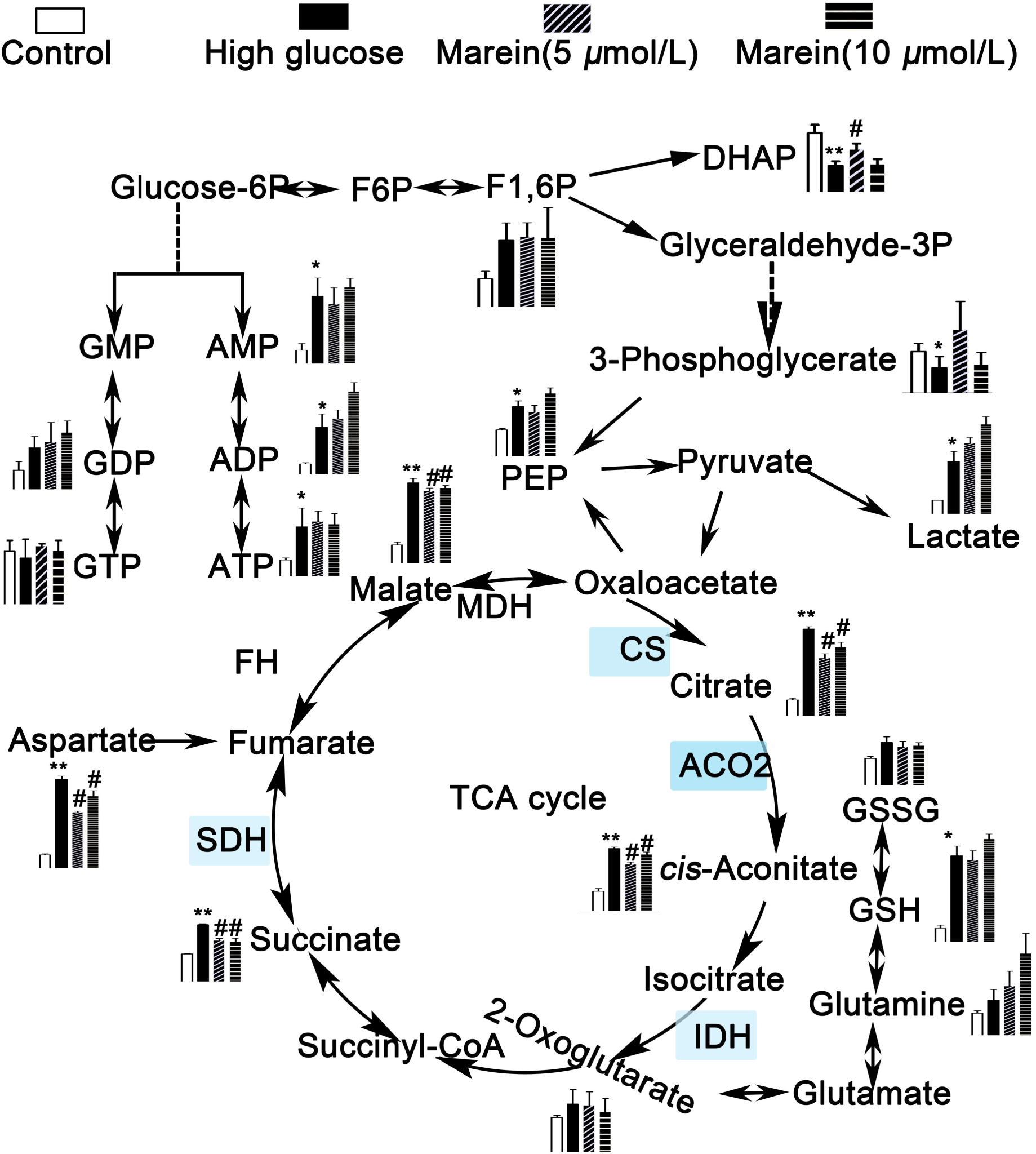
Metabolome pathway map of quantified metabolites, including components of the glycolytic or gluconeogenesis pathway and Krebs cycle in all groups. The red box shows the enzymes increased by high glucose treatment. \**P<0.05* vs the control group; ** *P<0.01* vs the control group;*** *P<0.001* vs the control group; # *P<0.05* vs 55 mmol/L glucose-treated group; ## *P<0.01* vs 55 mmol/L glucose-treated group. F6P, Fructose 6-phosphate; F1,6P, Fructose 1,6-bisphosphate; DHAP, Dihydroxyacetone phosphate; PEP, Phosphoenolpyruvate;

The high glucose condition clearly increased levels of other measured Krebs cycle intermediates compared with control group, including citrate (5.09-fold, p= 1.5×10^−7^), succinate (6.46-fold, p= 1.7×10^−4^), aconitate (3.11-fold, p= 5.2×10^−6^), and malate (4.08-fold, p= 8.8×10^−6^). 10 μmol/L and 5 μmol/L Marein partially decreased the effect of high glucose on these Krebs cycle intermediates.

### Marein reduces mRNA and protein expression of Krebs related enzymes SDHA, ACO2, IDH2 and CS in high-glucose-induced IR model cells

To evaluate the effect of Marein on Krebs cycle metabolism of human hepatocyte exposed to high glucose, the RNA and protein expression of Krebs related enzymes were detected with qPCR and western blot assays. As shown in figure 7A, treatment with 55 mmol/L glucose caused to an increase of some Krebs cycle mRNA levels, including *SDHA*, *ACO2*, *IDH2* and *CS*, whereas this effect was inhibited by Marein. There were no detectable differences in mRNA levels of *FH*, *MDH2* and *DLST*in the high-glucose-induced IR model cells. In accordance with the results of mRNA expression, the protein levels of SDHA, ACO2, IDH2 and CS were also increased in the treatment group with 55 mmol/L glucose, but FH, MDH2 and SDHB were not altered. Marein reduced the effect of high glucose on the protein expression of SDHA, ACO2, IDH2 and CS in a dose-dependent manner (Fig. 7B and 7C).

**Figure 7.**
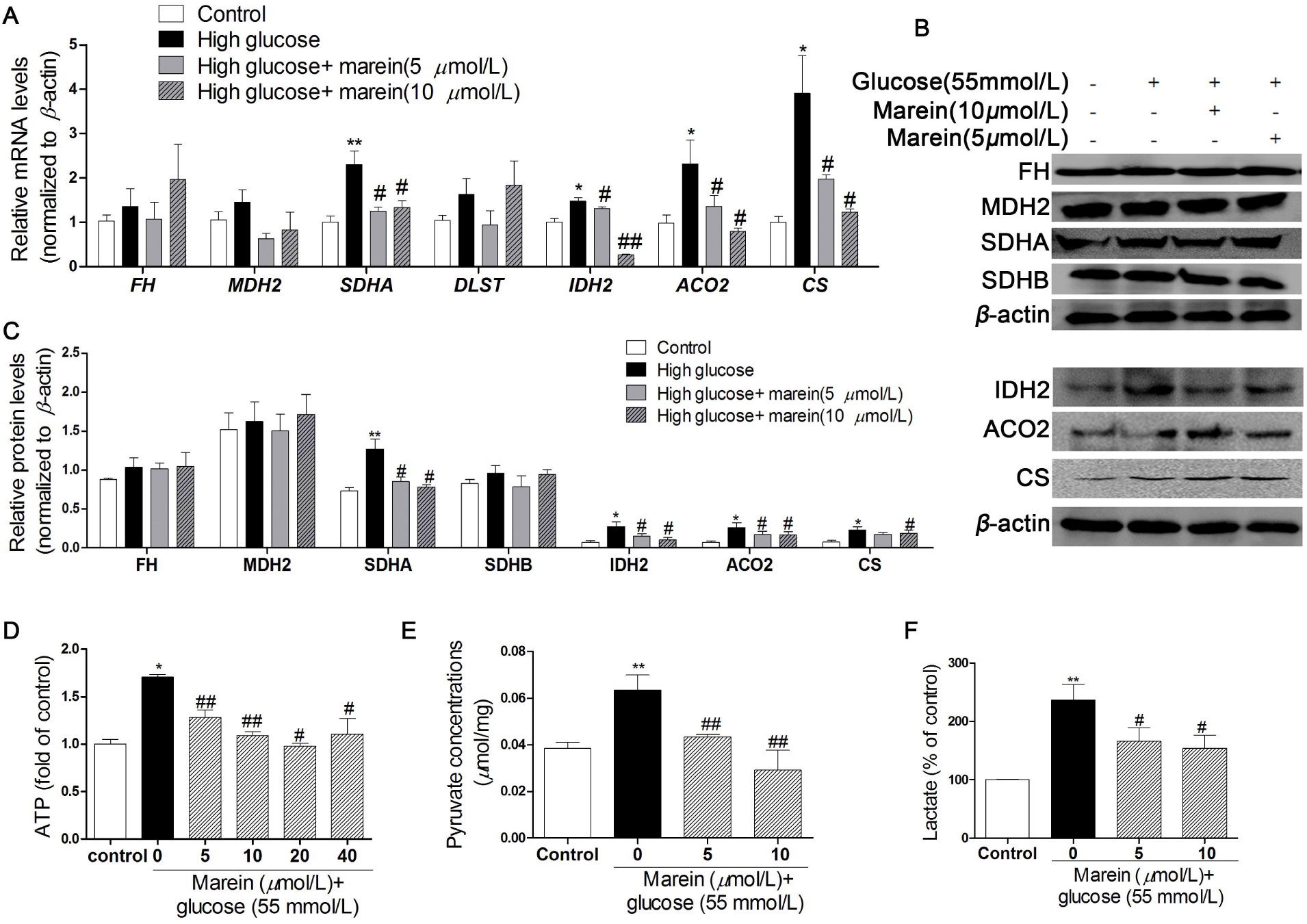
Marein inhibits the increase of mRNA expression and protein levels of Krebs cycle enzymes induced by high glucose in HepG2 cells. (A)Effect of Marein on mRNA expression of Krebs cycle genes. (B) Bands of representative experiments for FH, MDH2, SDHA, SDHB, IDH2, ACO2 and CS. (C) Densitometric quantification of Krebs cycle enzymes. (D) ATP levels expressed as a percent of control group are means ± SD of at least 4 different samples per condition. (E) Effect of Marein on pyruvate concentration. (F) Lactate concentrations expressed as a percent of control group are means ± SD of at least 4 different samples per condition.* *P<0.05* vs the control group; ** *P<0.01* vs the control group; # *P<0.05* vs 55 mmol/L glucose-treated group; ## *P<0.01* vs 55 mmol/L glucose-treated group.

### Marein reduces ATP levels, pyruvate and lactate concentration in high glucose-induced IR model cells

As exposure to high glucose will decrease glycogen synthesis, the effect of high glucose on gluconeogenesis was evaluated by detecting the pyruvate and lactate concentration and ATP levels. Under the IR condition, cells produced more ATP to supply energy due to a decline in glucose uptake. Treatment with high glucose also caused the elevation of pyruvate and lactate concentration, whereas this was inhibited by 10 and 5 μmol/L Marein (Fig.7D-F). This result indicates that increased gluconeogenesis plays a key role in the damage caused by high glucose in HepG2 cells, and that Marein can ameliorate that effect.

## Discussion

Previous study showed that the flower tea *C. tinctoria* increases insulin sensitivity and regulates hepatic metabolism in rats fed a high-fat diet^[22]^. In this study, it had been showed that AC strongly improves glucose and lipid homeostasis disrupted by HFD in rats. As the main components of AC, Marein protects HepG2 cells from high glucose-induced glucose metabolism disorder. Metabolomics analyses showed that Marein improves the multitude metabolic pathway including TCA cycle, gluconeogenesis and amino acid metabolism in HepG2 cells. Our results demonstrate that pretreatment of HepG2 cells with Marein significantly reversed high glucose-induced the changes of the key substrates and enzymes in gluconeogenesis and Krebs cycle.

The liver plays an important role in maintaining blood glucose concentration both by supplying glucose to the circulation via glycogenolysis and gluconeogenesis and by removing glucose from the circulation to increase glycogen synthesis ^[37]^. We found that glucose utilization and glycogen synthesis were significantly decreased but the activity of PEPCK and G6Pase, the key enzymes in the metabolic pathway of gluconeogenesis were obviously increased in cells treated with high glucose (Fig. 3). These results are in agreement with the previous findings that hepatic IR can be attributed mostly to decreased stimulation of glycogen synthesis by insulin while gluconeogenesis is abnormally enhanced due to the inefficient utility of glucose ^[39]^. Our data showing that pretreatment with Marein significantly reversed the high-glucose-induced decrease in glucose utilization and glycogen synthesis, increasing in the activity of PEPCK and G6Pase. Consistent with the protein expression of gluconeogenesis enzymes, the metabolites of glycolytic and gluconeogenesis pathway were significantly decreased in the high glucose group (Fig.. 3G), which could be reversed by pretreatment with Marein. In accordance with previous reports, pyruvate and lactate were clearly increased in the high glucose treatment cells ^[40, 41]^. These metabolites were the end product of glycolysis, thus their levels elevated in response to an increased glycolytic activity. This is different from the liver tissues with type 2 diabetes ^[42]^. One possible explanation for these results is that HepG2 cell is a kind of tumor cell line with a high rate of aerobic glycolysis, known as the Warburg effect, which is a hallmark of cancer cell glucose metabolism ^[43]^. Our results demonstrated that pretreatment with Marein before high glucose treatment significantly reduced the high glucose-induced increase in pyruvate and lactate generation and decrease in glycolytic activity in HepG2 cells. It is widely accepted that pyruvate is converted to acetyl-CoA and then enter the Krebs cycle in the mitochondria. Previous reports have demonstrated that the key enzyme activities of the Krebs cycle are significantly altered in type 2 diabetes ^[35, 44, 45]^.

In this study, mRNA expression and protein levels of these enzymes were measured, and it was found that treatment with high glucose caused an increase in mRNA and protein levels of some key enzymes including ACO2, IDH2, CS and SDHA, and affected upstream reactions of the Krebs cycle, findings which are in agreement with the previous findings that are mostly focused on the key enzymes activities of Krebs cycle ^[35, 44, 45]^. The high-glucose condition did not affect the enzymes downstream of the Krebs cycle, such as FH and MDH2. *SDHB* gene expression was not affected by high glucose, which may be because SDHB is not the catalyzing subunit (Fig. 7). However over-activation of these enzymes as induced by high glucose may be suppressed by pretreatment with Marein.

A recent report by Choi et al. ^[46]^ implicates glucolipotoxicity as a cause for impaired glucose metabolism, leading to a decrease in Krebs cycle intermediates that is consistent with our current findings that the levels of Krebs cycle intermediates, such as succinate, citrate, aconitate, and malate are significantly increased in the high glucose group. On the other hand, pretreatment with Marein before high glucose significantly increased the above Krebs cycle intermediates, which suggested that Marein could significantly increase the rate of the Krebs cycle and the accumulation of these intermediates, and thus reduce the toxicity of glucose. However, our study also found that mRNA and protein levels of Krebs cycle-related enzymes, such as SDHA, CS, ACO2 and IDH2, significantly increased in high glucose group and decreased in Marein pretreatment group, which may reverse the stress response to these substrates. The increase in these enzymes may degrade the related substrates.

In summary, we evaluated the effect of marein extracted from on IR. And then we have demonstrated that Marein attenuated the IR induced by high glucose in vitro. Our results show that Marein’s improvement of the IR effect can be attributed to it ameliorating numerous high glucose-induced processes such as decrease in glucose uptake and glycogen synthesis, inhibition the activity of hexokinase, perturbation of the glyconeogenesis and Krebs cycle homeostasis, and an increase in the levels of ATP, pyruvate and lactate. Meanwhile, Marein also improves glucose and lipid metabolism disorder disrupted by HFD in rat IR model. These findings suggest that Marein may have considerable potential for preventing high glucose-induced glucose metabolism disorder and IR.

## Acknowledgements

The work was supported by grants from the National Natural Science Foundation of China (8010907, 81271255 and 81274188), 973 Program (2013CB531200), the Program for Changjiang Scholars and Innovative Research Team (IRT1150). China HiTech 863 Program (2012AA020205, 2014AA020906), Instrumentation Program of Ministry of Science and Technology of China (2012YQ18011710), Young Scientist Grant from Peking Union Medical College (2012J22), CAMS Innovation Fund for Medical Sciences (CIFMS 2016-I2M-1-012).

## Author Contributions

Conceptualization: G,Xiao. Data curation: B,Jiang.

Formal analysis: LLe. Investigation: B,Jiang. Methodology: LLe.

Project administration: L, Xu K,Hu Resources: L, Xu.

Software: B,Jiang.

Supervision: G,Xiao.

Validation: G,Xiao.

Visualization: L, Xu K,Hu.

Writing – original draft: LLe B,Jiang. Writing – review & editing: LLe B,Jiang.

